# Tight control of the APP-Mint1 interaction in regulating amyloid production

**DOI:** 10.1101/2022.10.07.511363

**Authors:** Shawna M. Henry, Christian R. O. Bartling, Kristian Strømgaard, Uwe Beffert, Angela Ho

**Affiliations:** Department of Biology, Boston University, Boston, MA, USA; Department of Drug Design and Pharmacology, University of Copenhagen, Copenhagen, Denmark

**Keywords:** Mint1, APP, amyloid, Alzheimer’s disease

## Abstract

Generation of amyloid-beta (Aβ) peptides through the proteolytic processing of the amyloid precursor protein (APP) is one pathogenic event in Alzheimer’s disease (AD). APP is a type I transmembrane protein and endocytosis of APP mediated by the endocytic YENPTY sequence is a key step in Aβ generation. We and others have found that Mints, a family of cytosolic adaptor proteins, directly binds to the YENPTY motif of APP *via* phosphotyrosine binding (PTB) domain of Mints, facilitates APP trafficking and processing. We also show mutation of Tyr633 of Mint1 (Mint1^Y633A^) enhances APP binding and processing. Now, we created a low-affinity Mint1 mutant that targets two conserved residues, Tyr549 and Phe610 (Mint1^Y549A/F610A^), that reduced APP binding. Here, we investigate how perturbing the APP-Mint1 interaction alters APP and Mint1 cellular dynamics as well as Mint1’s interaction with its other binding partners. We show that Mint1^Y633A^ increased binding affinity specifically for APP and presenilin1, enhanced APP endocytosis, and Aβ secretion in primary neurons. Conversely, Mint1^Y549A/F610A^ exhibited reduced APP affinity and Aβ secretion. In fact, the effect of Mint1^Y549A/F610A^ on Aβ release was greater compared to knocking down all three Mint proteins, supporting targeting APP-Mint1 interaction as a potential AD therapeutic.

## 1. Introduction

Accumulation of amyloid-β (Aβ) peptides into insoluble plaques is a pathological hallmark of Alzheimer’s disease (AD). Aβ is produced by the sequential proteolytic processing of the amyloid precursor protein (APP) by β- and γ-secretases (Selkoe and Hardy, 2016). In neurons, APP accumulates in the trans-Golgi complex before trafficking to the plasma membrane, and undergoes a rapid turnover by α-secretase at the plasma membrane (non-amyloidogenic pathway). Alternatively, APP is internalized to endosomes where APP is cleaved by β-secretase, initiating the amyloidogenic pathway. The sorting signal that regulates APP endocytic processing required for Aβ generation is the conserved endocytic YENPTY sequence located in the cytoplasmic region of APP (Lai et al., 1995; Perez et al., 1999; Haass et al., 2012).

Mints (also known as APP binding family A, APBA) are a family of neuronal adaptor proteins that directly bind to the YENPTY motif of APP and regulate APP processing associated with AD (Ho et al., 2008). Mints are encoded by three distinct genes: neuron-specific Mint1 and Mint2 and the ubiquitously expressed Mint3 (Okamoto and Sudhof, 1997). Mints consist of a divergent N-terminus and a conserved C-terminus that encodes a phosphotyrosine binding (PTB) domain, an α-helical linker (ARM) domain, and two tandem PDZ domains. We and others have found the endocytic YENPTY motif of APP binds directly to the PTB domain of Mints, and is essential for regulating APP trafficking and processing (Borg et al., 1996; Chaufty et al., 2012; Sullivan et al., 2014). Through the crystal structure of Mint1, we found the ARM domain adjacent to the PTB domain folds back, and sterically hinders APP binding (Matos et al., 2012). A single point mutation in Y633A of Mint1 in the ARM domain has been shown to relieve autoinhibition, and increase APP binding. Conversely, recent studies found mutations in the Mint2 PTB domain (Y459A and F520A) led to reduced APP binding and decreased Aβ generation (Bartling et al., 2021).

Disrupting the APP-Mint interaction is a promising therapeutic avenue to selectively reduce Aβ production in AD. However, an outstanding question is whether targeting the APP-Mint interaction also interferes with other protein-protein interactions that may alter Mints function since Mints are multidomain proteins that bind to several AD and synaptic-associated proteins. Here we will examine two fulllength Mint1 mutants, Mint1^Y633A^ and Mint1^Y549A/F610A^ (which is analogous to Y459 and F520 in Mint2), to determine the binding specificity to APP and Mint1 interacting partners. In addition, we examined the cellular and molecular mechanistic effects of Mint1 mutants in neurons, and found Mint1^Y549A/F610A^ selectively decreased APP binding without interfering with Mint1 interacting proteins to decrease APP endocytosis and Aβ production in neurons.

## 2. Results

### 2.1 Mint1 mutants exhibit differential binding affinities to APP and Mint1 interacting partners

To assess Mint1’s ability to bind APP and other interacting partners, we produced two full-length GFP-tagged Mint1 mutants, GFP-Mint1^Y633A^ and GFP-Mint1^Y549A/F610A^ to compared with GFP-Mint1^WT^. We measured their ability to interact with the APP family of proteins including, APLP1 and APLP2 by cotransfecting with GFP-Mint1^WT^, GFP-Mint1^Y633A^, or GFP-Mint1^Y549A/F610A^ in HEK293T cells, and performing a co-immunoprecipitation assay. We immunoprecipitated with GFP antibody and immunoblotted for Mint1 and APP. GFP-Mint1^Y633A^ enhanced APP binding by 767% compared to GFP-Mint1^WT^ (lane 7 compared to 6, Fig. 1A). Meanwhile, GFP-Mint1^Y549A/F610A^ reduced APP binding by 82% compared to GFP-Mint1^WT^ (lane 8 compared to 6, Fig. 1A). Similarly, GFP-Mint1^Y633A^ enhanced Mint1 binding to APLP1 and APLP2 (lane 7 compared to 6, Fig. 1B, C). Conversely, GFP-Mint1^Y549A/F610A^ reduced binding to APLP1 and APLP2, which could be partially attributed to the decrease in APP, APLP1 and APLP2 protein stability when co-expressed with GFP-Mint1^Y549A/F610A^ (lane 8 compared to 6, Fig. 1B, C).

**Fig. 1.**
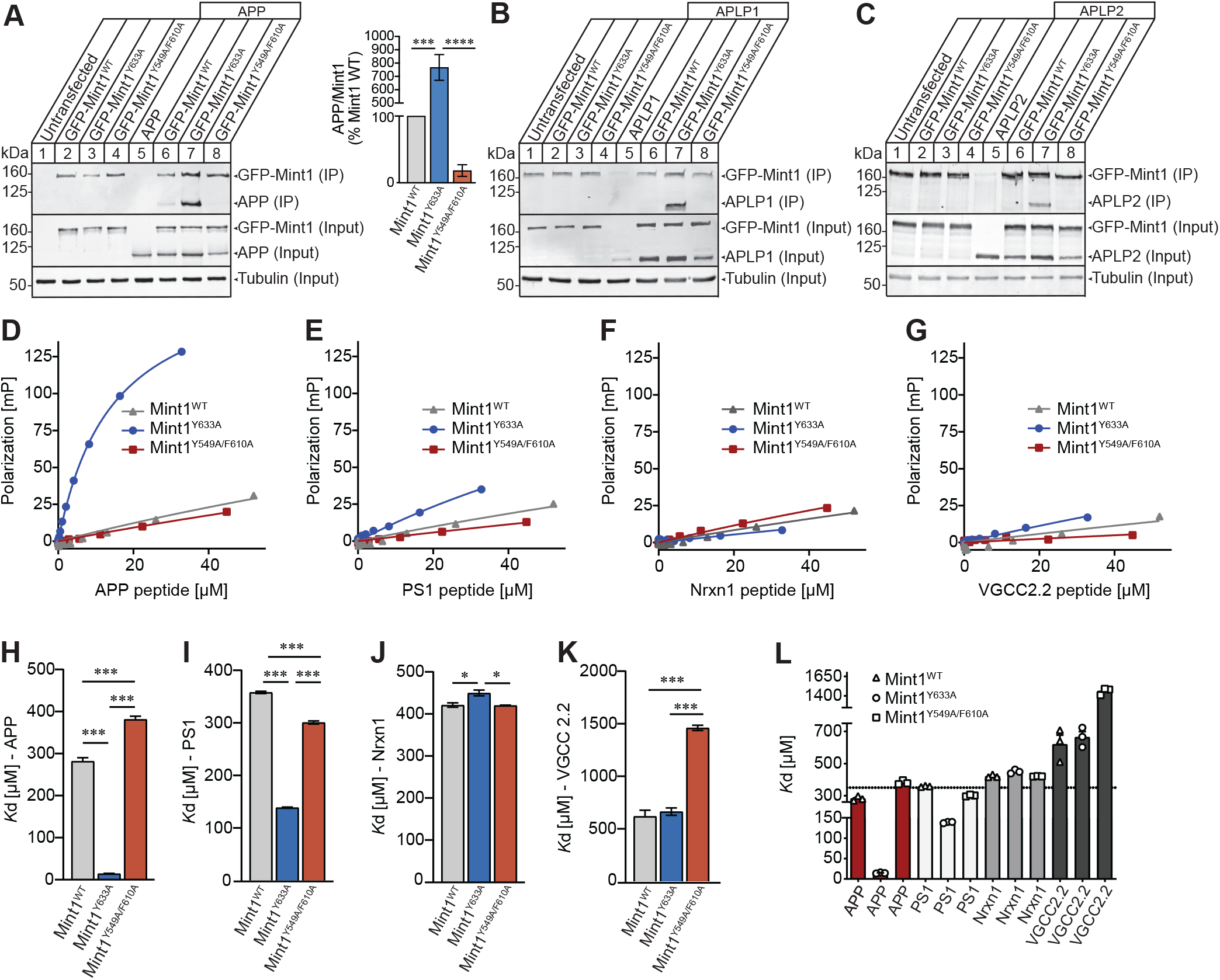
Biochemical analysis of Mint1 mutants with APP family of proteins and interacting partners. (A) HEK293T cells were co-transfected with APP and GFP-Mint1^WT^, GFP-Mint1^Y633A^, or GFP-Mint1^Y549A/F610A^ plasmids. Cell lysates were collected 48 hours post-transfection and immunoprecipitated (IP) with GFP antibody and immunoblotted for APP, GFP, and tubulin. The amount of immunoprecipitated APP was normalized to the amount of precipitated Mint1 and shown as percent Mint1^WT^ control. Data are expressed as the mean ± SEM (n = 4 independent experiments). Statistical significance was evaluated using one-way ANOVA with Sidak’s multiple comparison test, ***p = 0.00012 and **** p = 0.00009. (B-C) GFP-Mint1^Y633A^ enhanced Mint1 binding to APLP1 (B) and APLP2 (C). Conversely, GFP-Mint1^Y549A/F610A^ reduced binding to APLP1 and APLP2. (D-G) Fluorescence polarization (FP) saturation curves of the binding of APP peptide (D), PS1 (E), Nrxn1 (F), VGCC2.2 to recombinantly expressed Mint1 C-terminal constructs encompassing 453-839 amino acids that carried Mint1^WT^, Mint1^Y633A^, or Mint1^Y549A/F610A^ mutations. (H-K) Affinity fold-change to APP peptide (H), PS1 (I), Nrxn1 (J), VGCC2.2 (K) toward Mint1^WT^, Mint1^Y633A^, or Mint1^Y549A/F610A^ mutations obtained in a FP assay. (L) Summary comparison of fold change for APP and Mint1 interacting partners toward Mint1^WT^ and mutants (n *=* 1 independent experiment with 3 biological replicates). Statistical significance was evaluated using one-way ANOVA with Sidak’s multiple comparison test, *p < 0.05 and ***p < 0.0001.

To target the APP-Mint1 interaction therapeutically, the interference needs to be specific to APP, and not to other Mint1 interacting partners. To test this, we quantified the interaction of Mint1^WT^ and Mint1 mutants with known interacting partners such as presenilin 1 (PS1), cell adhesion molecule neurexin (Nrxn 1), and voltage-gated calcium channel 2.2 (VGCC 2.2) using fluorescence polarization (FP). We used recombinantly expressed Mint1 C-terminal constructs encompassing 453-839 amino acids that carried the Y549/F610A or Y633A mutations, and first tested the 17-mer APP C-terminal peptide that encompassed the endocytic YENPTY motif as a positive control. Based on the FP saturation curves of the APP peptide to Mint1 mutants, Mint1^Y633A^ showed enhanced binding affinity with Kd of 14.5 ± 0.74 μM, a 19-fold increase compared to Mint1^WT^ (K_d_ = 282 ± 8.3 μM) (Fig. 1D, H). Mint1^Y549A/F610A^ exhibited an even lower affinity to APP peptide (K_d_ = 382 ± 12.5 μM) compared to Mint1^WT^. Interestingly, we found Mint1^Y633A^ preferentially bound PS1 peptide with higher affinity (K_d_= 138.8 ± 1.02 μM), at least a 2-fold increase compared to Mint1^WT^(K_d_= 358 ± 2.0 μM), and Mint1^Y549A/F610A^ (K_d_= 301 ± 3.0 μM) (Fig. 1E, I). However, the binding affinity of Mint1^Y633A^ to PS1 peptide was not as strong compared to APP peptide. Based on the FP saturation curves for Nrxn1 and VGCC2.2 peptides to Mint1^WT^ and Mint1 mutants, the binding affinity was much lower with a Kd range of 420-664 μM (Fig. 1F, G, J, K). We observed the weakest affinity between Mint1^Y549A/F610A^ and the VGCC2.2 peptide (K_d_= 1461 ± 25.2 μM) compared to Mint1^WT^ (Kd= 619 ± 57.9 μM) (Fig. 1K). Overall, we found a single mutation in Y633A of Mint1 in the ARM domain specifically enhanced binding to both APP and PS1, suggesting Mint1^Y633A^ may increase APP endocytosis and facilitate γ-secretase dependent APP processing. Also, we found Y549 and F610 within Mint1 PTB domain are essential for the APP-Mint1 interaction because the Mint1^Y549A/F610A^ mutant exhibited a significant reduction in APP binding.

### 2.2 Mint1 mutants alters Golgi and APP co-localization in primary neurons

To determine whether the Mint1 mutants affect Mint1 localization in neurons, we cultured primary neurons from mice that lacked endogenous Mint1, and infected lentivirus that expressed GFP-Mint1^WT^, GFP-Mint1^Y633A^, or GFP-Mint1^Y549A/F610A^ at DIV 2. Since previous studies have shown that Mint proteins localize to the Golgi apparatus (Biederer et al., 2002), we immuno-labeled for Mint1 and the Golgi marker GM130 at DIV 10. Both GFP-Mint1^WT^ and GFP-Mint1^Y633A^ overlapped with GM130, with slightly more co-localization between GFP-Mint1^Y633A^ and GM130 compared to GFP-Mint^WT^ (Fig. 2A, B). Meanwhile, we observed a diffuse labeling of GFP-Mint1^Y549A/F610A^ and decrease co-localization with GM130. We also quantitatively assessed the Golgi morphology that has been linked to different cellular processes (Makhoul et al., 2019), with the Golgi defined as “condensed” when it appeared compact and nearly circular in shape, and “ribbon” when the Golgi is extended with multiple cisternae. We found neurons infected with GFP-Mint1^Y633A^ tend to have more “condensed” Golgi morphology compared to GFP-Mint1^WT^ and GFP-Mint1^Y549A/F610A^ (Fig. 2C).

**Fig. 2.**
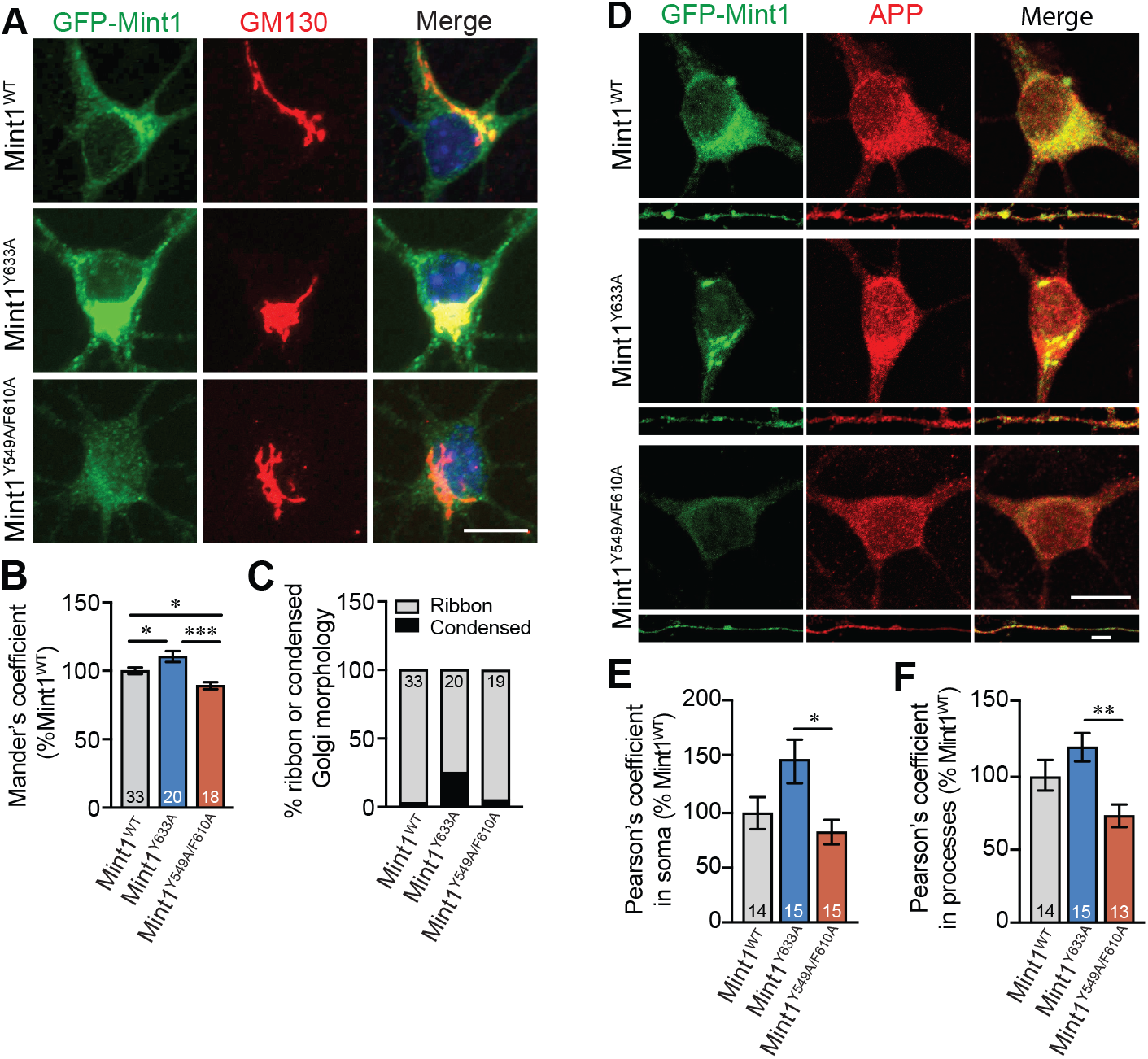
Cellular localization of Mint1 mutants in primary neurons. (A) Primary murine neurons that lacked endogenous Mint1 and infected with GFP-Mint1^WT^, GFP-Mint1^Y633A^, or GFP-Mint1^Y549A/F610A^ at 2 DIV and immunolabeled with GFP and cis-Golgi marker GM130 on 10 DIV. Representative images show GFP-Mint1 (green) and Golgi staining (red). Scale bar = 10 μm. (B) Co-localization of GFP-Mint1 with GM130 was quantified using Mander’s coefficient normalized to Mint1^WT^. Data are expressed as the mean ± SEM (n = 1 independent experiment, number of neurons analyzed is indicated at the bottom of each bar). Statistical significance was evaluated using one-way ANOVA with Sidak’s multiple comparison test, *p ≤ 0.05, ***p ≤ 0.001. (C) Percentage of neurons exhibiting a ribbon or condensed Golgi phenotype. Same neurons as analyzed in panel B. (D) Representative images show GFP-Mint1 (green) and APP staining (red). Scale bar for soma = 10 μm and processes = 5 μm. (E-F) Co-localization of GFP-Mint1 with APP was quantified in soma and processes using Pearson’s coefficient normalized to Mint1^WT^. Data are expressed as the mean ± SEM (n = 1 independent experiment, number at the bottom of each bar represents number of neurons analyzed). Statistical significance was evaluated using one-way ANOVA with Sidak’s multiple comparison test, *p ≤ 0.05, **p ≤ 0.01.

We next examined whether Mint1 mutants altered co-localization with APP by immuno-labeling for Mint1 and APP. Quantitative analysis of confocal images showed GFP-Mint1^WT^ and GFP-Mint1^Y633A^exhibited 43% and 38% higher co-localization with APP in both soma and processes compared to GFP-Mint1^Y549A/F610A^, respectively (Fig. 2D, E, F). These results suggest that the APP-Mint1 interaction is important for the localization of Mint1 to the Golgi which might be important to direct distinct cellular processes such as APP distribution across neurons.

### 2.3 Mint1 mutants alters APP endocytosis and Aβ production in primary neurons

Activity-dependent APP endocytosis is a critical step in Aβ production (Cirrito et al., 2005), and we have shown Mints are necessary for regulating APP endocytosis and Aβ production (Sullivan et al., 2014). To examine whether Mint1 mutants alter APP endocytosis and Aβ production, we cultured neurons from homozygous triple-floxed conditional Mint mice carrying the APPswe/PS1ΔE9 transgene that overproduce human Aβ (MTF^tg^). At DIV 2, neurons were infected with lentiviral *Cre* recombinase to knockdown all three Mint proteins, and rescued with either GFP-Mint1^WT^, GFP-Mint1^Y633A^, or GFP-Mint1^Y549A/F610A^lentivirus. Neuronal lysates collected at DIV 15 showed efficient knockdown for all three Mint proteins following *Cre* recombinase infection (lanes 3-10, Fig. 3A), whereas neurons infected with inactive *Cre*recombinase (Δ*Cre*) retain endogenous Mint expression (lanes 1-2, Fig. 3A). In addition, expression of GFP-Mint1^WT^, GFP-Mint1^Y633A^, or GFP-Mint1^Y549A/F610A^ lentivirus was comparable to endogenous Mint protein levels (lanes 5-10, Fig. 3A).

**Fig. 3.**
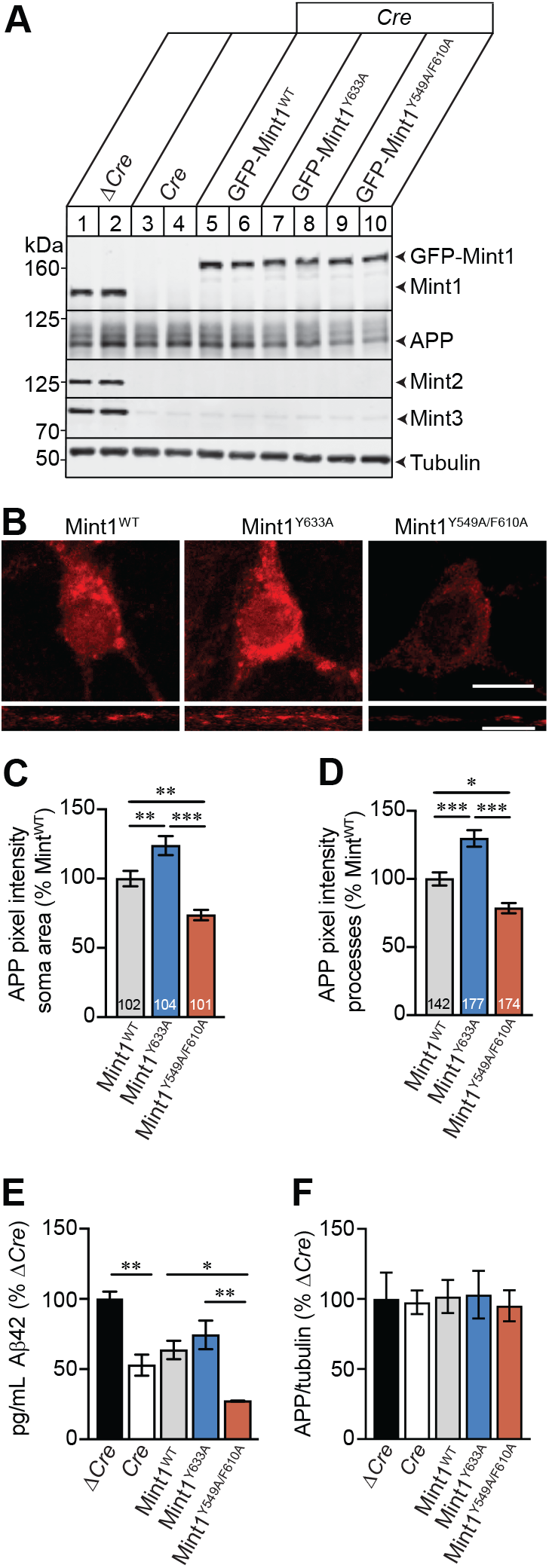
Mint1 mutants alter APP endocytosis and Aβ production in primary neurons. (A) Representative immunoblot analysis of lysates from Mint triple-floxed neurons carrying the APPswe/PS1ΔE9 transgene (MTF^tg^) that were infected with inactive lentiviral *Cre* recombinase (*ΔCre*) or active *Cre* recombinase to knockdown Mints 1-3 and rescued with GFP-Mint1^WT^, GFP-Mint1^Y633A^, or GFP-Mint1^Y549A/F610A^ lentivirus. Cell lysates immunoblotted for individual Mint proteins showed efficient knockdown of Mints 1-3 (lanes 3-10), and comparable expression of GFP-Mint1^WT^ (lanes 5-6), GFP-Mint1^Y633A^ (lanes 7-8), or GFP-Mint1^Y549A/F610A^ (lanes 9-10) lentivirus compared to endogenous Mint1 levels (lanes 1-2). Tubulin serves as a loading control. (B) Live endocytosis assay of primary MTF^tg^ neurons that were infected with *Cre* recombinase and rescue with GFP-Mint1^WT^, GFP-Mint1^Y633A^, or GFP-Mint1^Y549A/F610A^ lentivirus. Neurons were incubated with an extracellular N-terminal APP antibody and subsequently treated with 25 μM glutamate for 5 min at 37°C prior to immunostaining. Representative images showing internalized APP (red) in both the soma (top) and processes (bottom). Scale bars: soma = 10 μm; process = 5 μm. (C-D) Quantification of the amount of internalized APP using corrected total cell fluorescence (El-Sharkawey, 2016) in the neuronal soma (C) and processes (D), expressed as percent Mint1^WT^. Data are expressed as the mean ± SEM (*n*= 2 independent experiment, number at the bottom of each bar represents number of neurons analyzed). Statistical significance was evaluated using one-way ANOVA with Sidak’s multiple comparison test, *p ≤ 0.05, **p ≤ 0.01, ***p ≤ 0.001. (E) Aβ42 ELISA quantification of conditioned media from MFT^tg^ neurons. (F) Quantitative analyses of APP expression from MTF^tg^ lysates by Western blotting (panel A). Data were normalized to Δ*Cre* control, and expressed as the mean ± SEM (n = 1 independent experiment with 3 biological replicates). Statistical significance was evaluated using one-way ANOVA with Sidak’s multiple comparison test, *p ≤ 0.05, **p ≤ 0.01.

To stimulate activity-induced APP endocytosis, neurons were treated with glutamate for 5-min at DIV 15, and live-cell endocytosis for APP was performed. GFP-Mint1^Y633A^ induced a ~24% and 30% increase in internalized APP in the soma and processes compared to GFP-Mint1^WT^, respectively (Fig. 3B, C, D). In contrast, neurons infected with GFP-Mint1^Y549A/F610A^ exhibited a ~27% and 22% decrease in APP endocytosis in the soma and processes compared to GFP-Mint1^WT^, respectively. We next examined whether the changes in APP endocytosis reflects Aβproduction. We therefore, quantified Aβ42 levels released from primary neurons at DIV 15. As expected, Mint knockout neurons (*Cre*) exhibited a 44% decrease in Aβ42 production compared to neurons expressing Mints (Δ*Cre*) (Fig. 3E). Neurons infected with GFP-Mint1^WT^ exhibited a 32.7% decrease, a comparable decrease similar to Mint knockout neurons, which is likely due to the native autoinhibited state of Mint1 hindering its binding to APP. As GFP-Mint1^Y633A^ has been shown to bind APP at higher affinity and enhance APP endocytosis, infecting neurons with GFP-Mint1^Y633A^ caused Aβ production similar to neurons expressing all three endogenous Mints. Remarkably, GFP-Mint1^Y549A/F610A^ showed a robust 62.6% decrease in Aβ42 production compared to neurons expressing Mints. The decrease in Aβ production was not due to any changes in overall APP protein level changes across the different groups (Fig. 3F), suggesting the APP-Mint1 interaction is a critical factor in Aβ production.

## 3. Discussion

Here, we characterized two Mint1 mutants, Mint1^Y633A^ and Mint1^Y549A/F610A^ that bind to APP with high and low affinity, respectively. We previously found that a single point mutation in Mint1^Y633A^ relieved autoinhibition and enhances APP binding (Matos et al., 2012). Now, we showed Mint1^Y633A^ also facilitates PS1 binding compared to Mint1^WT^ supporting a mechanism whereby Mint1 promotes both APP and PS1 binding to enhance endocytic trafficking and processing of APP. In addition, we uncovered two conserved amino acids in the PTB domain of Mint1 Y549 and F610 (which is analogous to Y459 and F520 in Mint2), also reduced APP, APLP1 and APLP2 binding compared to Mint1^WT^ without affecting other Mint1 interacting partners such as PS1 and Nrxn1. Notably, Mint1^Y549A/F610A^ exhibited a two-fold decrease affinity (K_d_ = 1461 ± 25 μM) to VGCC2.2 as compared to Mint1 WT (K_d_ = 619 ± 58 μM). However, these are low-affinity interactions so this difference may not be physiologically pertinent.

Mints typically localize predominantly to the Golgi and synapse in primary neurons (Biederer et al., 2002; Okamoto et al., 2000). We found Mint1^Y633A^ exhibited increased colocalization with the Golgi. In contrast, Mint1^Y549A/F610A^ staining was more diffuse, with loss of colocalization with GM130, suggesting Mint1’s Golgi localization may be dependent on its interaction with APP. This is supported by previous work which showed Mint3 was recruited to the Golgi in an APP-dependent manner in Hela cells (Caster and Kahn, 2013). Further studies in *Drosophila* showed that Mints function at the Golgi to control polarized trafficking of axonal membrane proteins including APP (Gross et al., 2013). In fact, Mints have been shown to bind to ADP ribosylation factor, which function at the Golgi to facilitate the sorting of membrane proteins for transport (Hill et al., 2003). Considering Mint1^Y549A/F610A^ loss its colocalization with GM130, this may perturb APP trafficking from the Golgi. Together, these experiments suggest that Mint’s neuronal localization is dependent on its protein-protein interactions, especially with APP.

We also found Mint1^Y633A^ expressing neurons showed a greater number of neurons with a condensed Golgi morphology compared to Mint1^WT^ and Mint1^Y459A/F520A^ mutant. Since the Golgi has been implicated as a site for amyloidogenic APP processing (Fourriere and Gleeson, 2021), and alterations in Golgi morphology are a preclinical feature that occurs before neurodegeneration associated with AD (Dal Canto, 1996; Stieber et al., 1996), it is possible that Mints could also function in APP processing at the Golgi. Neurons expressing Mint1^Y549A/F610A^ showed a decrease in APP endocytosis and Aβ release when compared to both Mint1^WT^ and Mint1^Y633A^. The effect of Mint1^Y549A/F610A^ on Aβ production was greater compared to knocking down all three Mint proteins suggesting that targeting the APP-Mint1 protein-protein interaction may provide an alternative strategy without disrupting an adaptor protein with such a complex proteinprotein interaction network.

## 4. Conclusion

Here, we examined the biochemical and cellular dynamics of the APP-Mint1 interaction using two Mint1 mutants that bind APP high affinity (Mint1^Y633A^) or low affinity (Mint1^Y549A/F610A^). These Mint1 mutants exhibited profound alterations in cellular localization, APP endocytosis, and Aβ production,supporting the facilitative role of Mint1 in mediating APP trafficking and processing.

## 5. Experimental procedure

### 5.1 Plasmids

pEGFP-Mint1 was derived from *Rattus norvegicus* cDNA (NCBI NP_113967.1), and the pCMV5-APP695 construct was derived from *Homo sapiens* (NCBI NP_958817.1). To generate the rat pEGFP-Mint1^Y633A^ mutation, site-directed mutagenesis was performed using QuikChange II Site-Directed Mutagenesis Kit (Agilent Technologies) with primer set:SMH1607 forward GAAGACCTGAGCCAGAAGGAG*GCA*AGCGACCTGCTCAACACCCAG and SMH1608 reverse CTGGGTGTTGAGCAGGTCGCT*TGC*CTCCTTCTGGCTCAGGTCTTC. To generate the rat pEGFP-Mint1^Y549A/F610A^ mutation, we first mutated F610A mutation using QuikChange II Site-Directed Mutagenesis Kit with primer set:SMH1721 forward, CAGTCCATCGGGCAGGCC*GC*CAGCGTTGCATACCAGGAG and SMG1722 reverse CTCCTGGTATGCAACGCTG*GC*GGCCTGCCCGATGGACTG. We then consecutively mutated the Y549A mutation using the Q5 Site-Directed Mutagenesis Kit (New England Biolabs) with primer set: SMH1747 forward, GACCATTTCC*GC*CATCGCAGACATTG and SMH1748 reverse CTCAGAGGGTGGTCCATC. The subsequent pEGFP plasmids were subcloned into the lentiviral pFUW vector.

### 5.2 Co-Immunoprecipitation in HEK293T cells

HEK293T (ATCC CRL-3216) cells were maintained in Dulbecco’s Modified Eagle’s Medium (DMEM; Gibco) supplemented with 10% fetal bovine serum (FBS; Atlanta Biologicals) and 1% Penicillin/Streptomycin (Gibco) at 37 °C in a 5%CO2 incubator. At 60% confluency, HEK293T cells were transfected using FuGENE®6 (Roche) following the manufacturer’s protocol. HEK293T cells were lysed in immunoprecipitation buffer containing: 10 mM Tris HCl pH 8.0, 150 mM NaCl, 1 mM EDTA pH 8.0, and 1% Triton X-100, supplemented with protease and phosphatase inhibitors 48 h following transfection. The cell lysate was sonicated and rotated for 30 min at 4°C and cleared with a 21,130*g* spin for 5 min. Protein extracts were incubated for 2 h with rotation at 4°C with the precipitating antibody, followed by overnight incubation at 4°C with either 10 μl of pre-equilibrated protein A or G Ultralink beads (Thermo Scientific). The resin was washed several times with immunoprecipitation buffer spinning at 700 *g* for 30 sec, and precipitated proteins were eluted by boiling for 10 min in reducing 2X sample reducing buffer and resolved by SDS-PAGE.

### 5.3 Western blotting

Lysates were separated on SDS-PAGE gels and transferred to a nitrocellulose membrane (GE Healthcare). Membranes were blocked for 1 h at room temperature with either 5% milk in Tris-buffered saline plus Tween (TBST) or Odyssey Blocking Buffer in phosphate-buffered saline (PBS) (LI-COR Biosciences) and incubated with primary antibodies overnight at 4°C. Following primary antibody incubation, membranes were washed three times with either TBST or PBS and incubated in the appropriate secondary antibody conjugated to either IRDye^®^680RD- or IRDye^®^800CW (LI-COR Biosciences) atTubulin serves as a loading control 1:20,000 in either 5%milk in TBST or Odyssey Blocking Buffer in PBS (LI-COR Biosciences) for 1 h at room temperature. Membranes were washed three times in the appropriate buffer (TBST or PBS) and imaged on the Odyssey®CLX Imaging System (LI-COR Biosciences).

### 5.4 Primary murine neurons

Primary neuronal cultures were prepared using either Mint mouse line that is *Mint1* and *Mint3* double knockout with floxed *Mint2* (*Mint1^-/-^;fMint2/fMint2;Mint3^-/-^*) or Mint triple-floxed mouse line (MTF^tg^) that is homozygous floxed for all three *Mint* genes (*fMint 1fMint1*; *JMint2fMint2*; *JMint3fMint3*) carrying the APPswe/PresenilinlΔexon9 transgene that overproduces human Aβ(Jackson Laboratory stock #004462) (Ho et al., 2008). Newborn pups of either sex were collected at postnatal day 0 (P0) and their dissociated brain tissue was trypsinized for 10 min at 37°C, triturated, and plated onto either Matrigel (Corning) coated glass coverslips or Matrigel (Corning) coated wells. Neuronal cultures were maintained in a 37°C humidified incubator with 5% CO_2_. To prevent glial proliferation, 4 μM Cytosine beta-D-arabinofuranoside hyrdrochloride (Ara-C; Sigma) was supplemented into the media on either 1 day *in vitro*(DIV) (hippocampal) or 2 DIV (cortical).

### 5.5 Lentiviral infection of mouse neurons

Recombinant lentiviruses were produced by transfecting HEK293T cells using FuGENE^®^6 (Roche) with plasmids encoding viral enzymes and envelope proteins essential for packing of viral particles (pRSV-EV, pMDLg/pRRE, and pCMV-VSVG) with the addition of a shuttle vector encoding the gene of interest (pFUW). The media was changed to neuronal growth media 24 h after transfection and the conditioned media was collected, spun at 1,000 x *g* for 10 min at 4°C, and filtered using a 0.45 μM filter (GE Healthcare) 48 h after transfection. Neurons were infected with lentiviral *Cre* recombinase at day of plating, while all other lentiviruses were infected at 2 DIV. Neurons were maintained in the same media/lentivirus mixture until analysis.

### 5.6 Immunocytochemistry and image analysis

Primary hippocampal cultures were prepared from newborn mice and plated on Matrigel-treated (Corning) glass coverslips. Neurons were fixed with pre-warmed 4% paraformaldehyde (Electron Microscopy Sciences) at room temperature for 8 min, washed 3 times with PBS (Invitrogen), and permeabilized and blocked in 10% goat serum (Invitrogen) and 0.1% saponin (Sigma) in PBS for 1 h at room temperature. Neurons were incubated with primary antibodies in blocking buffer (10% goat serum in PBS) overnight at 4°C. Neurons were washed 3 times with PBS and incubated with a secondary antibody conjugated to an Alexa Fluor®(Invitrogen) in blocking buffer (10% goat serum in PBS) for 1 h at room temperature. Following PBS washes, the coverslips were mounted using ProLong Gold Antifade Mountant with DAPI (Invitrogen) and imaged using Z-stacks with a Carl Zeiss LSM 700 confocal microscope at 63X magnification. Corrected total cell fluorescence (CTCF) of maximum intensity projections was acquired using FIJI (National Institute of Health (NIH)). Co-localization was quantified using IMARIS image analysis software (Oxford Instruments).

### 5.7 Live-cell APP endocytosis assay

Primary hippocampal cultures were prepared and plated onto Matrigel-coated glass coverslips. Neurons were infected with 20% *Cre* recombinase lentivirus at day of plating and 5% Mint1 lentivirus on 2 DIV. At 15 DIV, live hippocampal cultures were incubated with an N-terminal APP antibody (mouse anti-APP 22C11, 1:500; EMD Millipore) diluted in conditioned neuronal media for 15 min at room temperature. Neurons were washed with conditioned neuronal media twice to remove any unbound antibodies. Next, neurons were treated with 25 μM glutamate for 15 min at 37°C to allow internalization, and fixed with prewarmed 4% paraformaldehyde (Electron Microscopy Sciences). Any APP antibodies remaining on the cell surface were quenched with a non-florescent goat anti-mouse secondary antibody conjugated to horseradish peroxidase (Cell Signaling) diluted in 10% goat serum (Invitrogen) in PBS for 1 h at room temperature. Neurons were washed twice with PBS and permeabilized with 10% goat serum supplemented with 0.3% Triton-X-100 in PBS for 10 min, and blocked in 10% goat serum in PBS for 35 min. Neurons were then incubated with goat anti-mouse Alexa Fluor®546 1:500 (Fisher Scientific) in 10% goat serum in PBS for 1 h. Following three PBS washes, the coverslips were mounted using ProLong Gold Anti-fade Mountant with DAPI (Invitrogen) and imaged with a Carl Zeiss LSM 700 confocal microscope.

### 5.8 Aβ42 ELISA

To perform an Aβ42 ELISA, conditioned neuronal media was diluted and handled according to the protocol of the Human Aβ42 Ultrasensitive ELISA Kit (Invitrogen) followed by colorimetric readout at 450 nm.

### 5.9 Fluorescence polarization

The binding affinities were determined in a flat bottom black 384-well plate (Corning Life Science) using a Safire plate reader (Tecan). All experiments were conducted in 150 or 500 mM NaCl, 25 mM HEPES, 1% bovine serum albumin (BSA), pH 7.4 at 25°C. Fluorescence was measured at excitation/emission wavelength at 530/580 nm. The instrumental Z-factor was adjusted to maximum fluorescence and the G-factor was calibrated to give an initial milli-polarization at 20. Fluorescence polarization assays were performed as saturation experiments using TAMRA-APP17-mer [(TAMRA)-NNG-QNGYENPTYKFFEQMQN], TAMRA-PS1-10mer [(TAMRA)-NNG-QLAFHQFYI]; TAMRA-Nrxn1-10mer [(TAMRA)-NNG-KKNKDKEYYV]; TAMRA-VGCC2.2-20mer [(TAMRA)-NNG-LSSGGRARHSYHHPDQDHWC] at a concentration of 50 nM.

### 5.10 Statistical analysis

All statistical analyses were performed using Prism 9 software (GraphPad). To determine statistical significance, we used one-way analysis of variance (ANOVA) coupled with either Sidak’s or Dunnett’s multiple comparisons test (specified on each figure). All graphs depict mean ±standard error of the mean (SEM).

## Declaration of Competing Interest

The authors declare no competing financial interests.

## Funding

This work was supported by National Institute of Health Grants R01 AG044499 (A. H.); R21 AG072433 (A.H.); and the Harold and Margaret Southerland Alzheimer’s Research Fund.

## CRediT authorship contribution statement

**Shawna M. Henry:** Conceptualization, Methodology, Data curation; Formal analysis; Investigation; Validation; Visualization; Writing - original draft; Writing - review & editing. **Christian R. O. Bartling**: Methodology, Data curation; Formal analysis; Investigation; Validation. **Kristian Strømgaard**: Project administration; Resources; Supervision. **Uwe Beffert:** Supervision; Writing - review & editing. **Angela Ho**: Conceptualization; Funding acquisition; Project administration; Resources; Supervision; Writing - original draft; Writing - review & editing.

## Notes

### Competing Interest Statement

The authors have declared no competing interest.

